# Data-driven Cognitive Clusters in Persistent Developmental Dyslexia

**DOI:** 10.1101/2025.11.24.690327

**Authors:** Pedro Pinheiro-Chagas, Margo Kersey, Nilgoun Bahar, Sarah Inkelis, Julian Siebert, Jet M. J. Vonk, Rian Bogley, Ellie Carpenter, Dolce Vita Martin-Moreno, Jessica De Leon, Boon Lead Tee, Maria Luisa Mandelli, Christa Pereira, Zachary Miller, Maria Luisa Gorno-Tempini

**Author notes:** These authors equally contributed to this work (Pedro Pinheiro-Chagas, Margo Kersey).

## Abstract

Developmental dyslexia is a neurodevelopmental condition characterized by persistent reading difficulties despite adequate intelligence and educational opportunities. The cognitive underpinnings of dyslexia remain incompletely understood. Although multiple causal pathways have been proposed, there is a gap in studies that can reliably identify dyslexia subtypes based on comprehensive evaluations that go beyond measures of reading and phonology. We employed one of the largest detailed samples of children with developmental dyslexia resistant to intervention, consisting of 147 extensively phenotyped school-age children aged 7-14 years who were recruited from specialized schools for students with learning differences. We implemented data-driven analyses leveraging a comprehensive neuropsychological assessment to delineate cognitive subdomains and characterize the heterogeneity in the sample. Hierarchical clustering of 24 cognitive measures aided in constructing a framework to subsequently interpret emerging clusters. Four clusters emerged, predominantly associated with: Processing Speed & Executive Function, Visuospatial & Mathematical Reasoning, Phonological Manipulation, and Verbal Short-term Memory & Word Retrieval. Subsequent latent profile analysis identified two distinct dyslexia profiles: one characterized by marked difficulties in executive functioning and processing speed, and another displaying impairments primarily in verbal short-term memory and word retrieval relative to the other profile. Importantly, both profiles exhibited comparable severity of persistent reading difficulties despite their divergent cognitive profiles. These findings show that phenotypically similar reading impairments can arise from distinct underlying cognitive factors, emphasizing the role of comprehensive neuropsychological evaluation in developing tailored interventions for affected children.

## 1. Introduction

Developmental dyslexia is a neurodevelopmental condition marked by difficulties in learning to read despite sufficient intelligence and educational opportunities (Lyon et al., 2003; Stanovich, 1988). Prevalence estimates vary, with figures ranging from 3% to 17.5% (Wagner et al., 2020). Dyslexia can negatively affect emotional wellbeing, academic attainment, and later occupational prospects (Al-Lamki, 2012; Georgiou et al., 2024; Wilmot et al., 2023).

Reading is a complex skill involving auditory and phonological processing, visual and orthographic knowledge, working memory, and executive function (Bowers & Wolf, 1993; Patael et al., 2018; Snowling, 2008; Wolf & Bowers, 1999; Wolf et al., 2024). Weaknesses in any of these domains can impact reading. Since these processes interact, children with dyslexia present with diverse profiles, complicating both research and clinical practice (Lyon et al., 2003; Peterson & Pennington, 2012). Therefore, a better understanding of individual strengths and weaknesses is crucial for developing effective interventions (Lorusso et al., 2024; Nukari et al., 2020; Snowling & Hulme, 2012b).

Much debate has centered on the diagnosis and theoretical underpinnings of dyslexia. Early accounts, such as the phonological deficit hypothesis, emphasized difficulties in phonological awareness as the core deficit (Bradley & Bryant, 1983; Stanovich & Siegel, 1994). Later, the double-deficit hypothesis proposed naming speed (efficiency of accessing phonological information, typically assessed using rapid automatic naming tests) as another core difficulty. Accordingly, deficits in phonology, naming speed, or both contribute to dyslexia, with the combined profile leading to more severe difficulties (Wolf & Bowers, 1999).

While informative, these models do not capture the full range of profiles observed. More recent perspectives, including the multifactorial hypothesis, posit that dyslexia arises from the interplay of multiple cognitive, linguistic, and environmental risk factors (Catts & Petscher, 2022; Wolf et al., 2024).

Consistent with this view, dyslexia often co-occurs with other neurodevelopmental conditions. For example, 18–42% of children with dyslexia also meet criteria for attention-deficit/hyperactivity disorder (ADHD; Germanò et al., 2010), and up to 50% of children with either dyslexia or ADHD show impaired executive functioning (Al Dahhan et al., 2022). Dyslexia also frequently co-occurs with developmental language disorder (Adlof & Hogan, 2018; Snowling et al., 2019), in which reading comprehension is further impaired by broader language difficulties (Snowling et al., 2020). Intervention studies further reinforce the importance of heterogeneity. Intensive phonics-based programs reliably improve decoding and phonological awareness (Shaywitz et al., 2004; Temple et al., 2003), yet many children with dyslexia continue to experience persistent reading difficulties (Pennington, 2006; Snowling & Hulme, 2012b; Wolf & Bowers, 1999). Thus, while phonological training is necessary, it is often not sufficient to remediate all aspects contributing to reading difficulty.

Despite these considerations, many studies and clinical protocols continue to focus primarily on phonological processing and naming speed deficits, leaving other domains, such as executive functioning, verbal short-term memory, or visuospatial skills often unaddressed (Wokuri et al., 2023; Zoubrinetzky et al., 2014). Past work has also been limited by small sample sizes and inconsistent criteria, with studies either using IQ-reading discrepancy definitions (O’Brien et al., 2012; Wolf et al., 2024) or involving small cohorts (Heim et al., 2008; Ramus et al., 2003). Data-driven approaches offer a methodological solution, enabling the move beyond theory-driven categorizations to uncover patterns of impairment that might otherwise remain unexplored.

The present study addresses these gaps by leveraging a large, well-characterized cohort of children with dyslexia who completed an extensive neuropsychological battery spanning more than 30 standardized measures across reading, phonological processing, executive functioning, processing speed, working memory, and verbal short-term memory. We adopted a data-driven analytic pipeline, beginning with hierarchical clustering of task-level measures to reduce dimensionality, followed by latent profile analysis (LPA) to detect subgroups with similar cognitive profiles. We hypothesized this strategy would uncover distinct profiles that capture variability in core reading and phonological skills while systematically integrating higher-order cognitive domains, thereby advancing understanding of dyslexia’s heterogeneity and potentially providing a foundation for more individualized assessment and intervention strategies.

## 2. Materials & Methods

### 2.1 Participants

The sample of this study was selected from a large dataset of children who were referred to the University of California, San Francisco (UCSF) Dyslexia Center either with suspected or a formal diagnosis of dyslexia. Most of these children were recruited through the UCSF Dyslexia Center’s partner schools, Charles Armstrong School and Chartwell School, which are both private institutions located in the San Francisco Bay Area dedicated to supporting students with language-based learning differences, particularly dyslexia. All children participated in extensive multidisciplinary evaluation at the UCSF Dyslexia Center involving neurologists, neuropsychologists, psychiatrists, speech-language pathologists, and academic specialists. Diagnostic decisions were made during consensus conferences, in which the clinical team reviewed all available information, including developmental and family history, test results, behavioral observations, and interviews with parents and teachers (explained in more detail in sections 2.2–2.4. Participants were recruited through a specialized clinical referral pathway to ensure diagnostic homogeneity. Given the challenges inherent in recruiting this specific neurodevelopmental population, the sample size was maximized via retrospective analysis of comprehensive neuropsychological data. To ensure the stability of our data-driven analyses, only participants with >90% completion of the standardized assessment battery were retained. The final cohort consisted of 147 native English-speaking children with confirmed dyslexia (*M* ± *SD* = 10.6 years ± 1.7, 69 female, 20 non-right-handed). This sample size offered sufficient statistical power to robustly identify phenotypic subgroups using our multivariate statistical modeling.

**Table 1.**
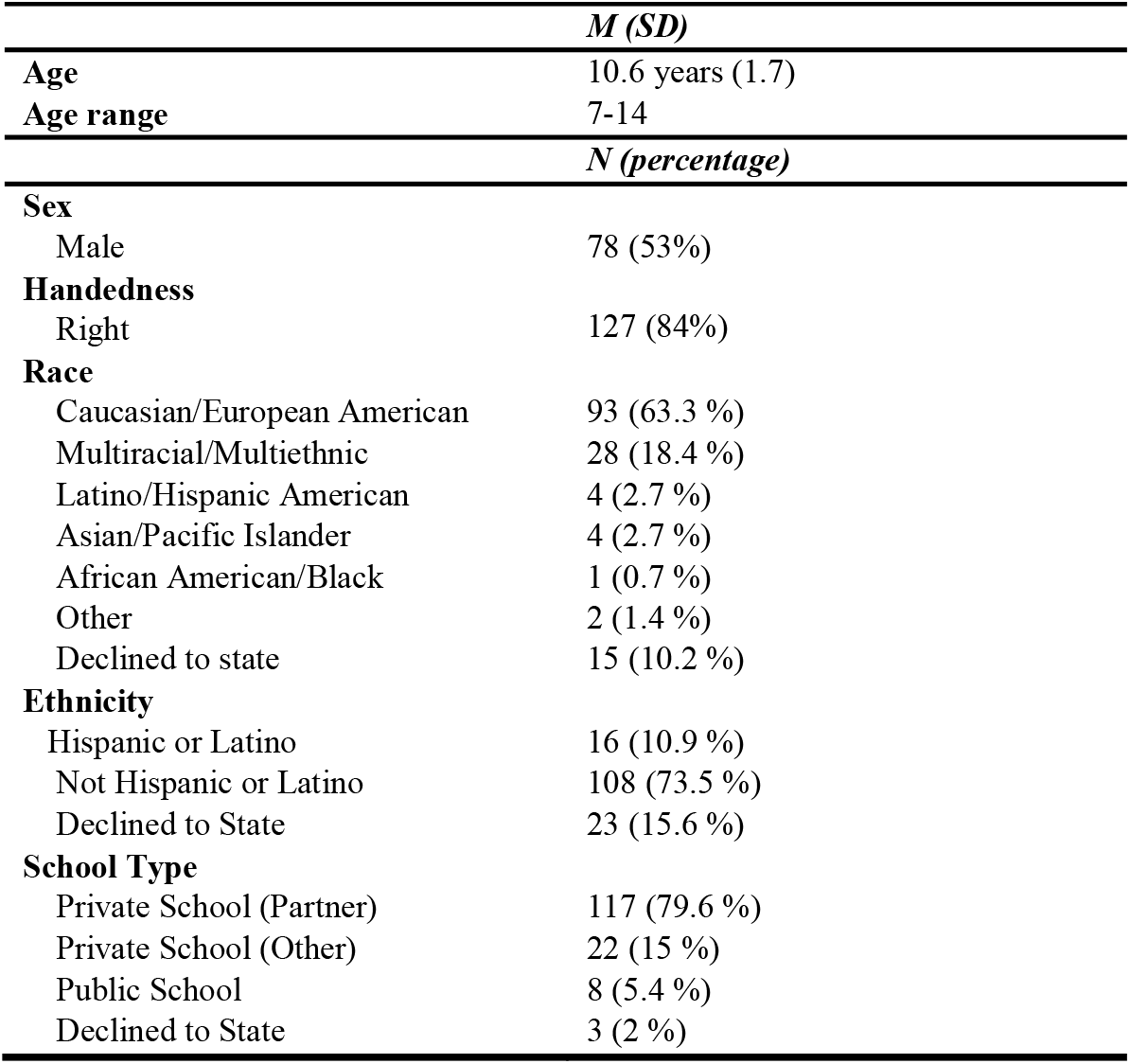
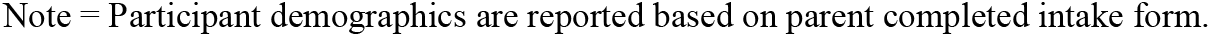
Participant Demographics Summary (N = 147)

|  | *M (SD)* |
| --- | --- |
| **Age** | 10.6 years (1.7) |
| **Age range** | 7-14 |
|  | *N (percentage)* |
| **Sex** |  |
| Male | 78 (53%) |
| **Handedness** |  |
| Right | 127 (84%) |
| **Race** |  |
| Caucasian/European American | 93 (63.3 %) |
| Multiracial/Multiethnic | 28 (18.4 %) |
| Latino/Hispanic American | 4 (2.7 %) |
| Asian/Pacific Islander | 4 (2.7 %) |
| African American/Black | 1 (0.7 %) |
| Other | 2 (1.4 %) |
| Declined to state | 15 (10.2 %) |
| **Ethnicity** |  |
| Hispanic or Latino | 16 (10.9 %) |
| Not Hispanic or Latino | 108 (73.5 %) |
| Declined to State | 23 (15.6 %) |
| **School Type** |  |
| Private School (Partner) | 117 (79.6 %) |
| Private School (Other) | 22 (15 %) |
| Public School | 8 (5.4 %) |
| Declined to State | 3 (2 %) |

Note = Participant demographics are reported based on parent completed intake form.

### 2.2 Test Battery & Questionnaires

The study was approved by the UCSF Institutional Review Board and conducted in accordance with the Declaration of Helsinki (World Medical Association, 2013). Informed written consent was obtained from parents or guardians, and assent was obtained from all participants. Families received a small monetary reimbursement for their time, as well as a conference and a detailed report summarizing their child’s results and providing educational recommendations.

The neuropsychological test battery encompassed tests of non-verbal reasoning, receptive and expressive language, phonological manipulation, phonological short-term memory, processing speed, short-term and working memory, learning and long-term memory, rapid naming, semantic fluency, visuospatial skills, visuomotor integration, and mathematical calculations (see Table 2 in Results for details). Assessment of reading included measures of single word/pseudoword reading from Test of Word Reading Efficiency– Second Edition (TOWRE-2; Torgesen et al., 2012) and Woodcock-Johnson, 4^th^ Edition—Tests of Achievement (WJ-IV; Schrank et al., 2014), paragraph reading from Gray Oral Reading Test–Fifth Edition (GORT-5; Wiederholt & Bryant, 2012), and single word/pseudoword spelling from WJ-IV (Schrank et al., 2014). Non-verbal intelligence was assessed using the Matrix Reasoning subtest of the Wechsler Abbreviated Scale of Intelligence (WASI; Wechsler, 1999) and receptive vocabulary was assessed using the Receptive One-Word Picture Vocabulary Test–Fourth Edition (ROWPVT-4; Martin & Brownell, 2011).

**Table 2.** Neuropsychological assessment results from 147 children with dyslexia.

|  | *M (SD)*<br>(%ile) | Range | *N* Missing Scores |
| --- | --- | --- | --- |
| **Reasoning Estimates** |  |  |  |
| WASI Matrix Reasoning | 61.2 (23.7) | 9-98 |  |
| ROWPVT-4 | 70.5 (23.4) | 13-99.8 |  |
| **Single Word Reading** |  |  |  |
| TOWRE-2 Sight Word Efficiency | 14.0 (16.8) | 0.1-70 |  |
| TOWRE-2 Phonemic Decoding Efficiency | 11.5 (14.2) | 0.1-58 |  |
| WJ-IV Letter Word Identification | 21.9 (20.6) | 0.1-88 |  |
| WJ-IV Word Attack | 30.4 (22.0) | 0.3-92 |  |
| **Oral Paragraph Reading** |  |  |  |
| GORT-5 Rate | 14.9 (14.8) | 0.1-91 | 1 |
| GORT-5 Accuracy | 8.7 (9.0) | 0.1-50 | 1 |
| GORT-5 Comprehension | 20.2 (17.3) | 0.4-75 | 1 |
| **Single Word Spelling** |  |  |  |
| WJ-IV Spelling | 12.0 (13.4) | 0.02-61 | 1 |
| WJ-IV Spelling of Sounds | 31.7 (19.3) | 0.1-86 | 1 |
| **Phonological Manipulation** |  |  |  |
| WJ-IV Segmentation | 60.5 (24.8) | 0.1-99 | 2 |
| WJ-IV Sound Blending | 56.7 (24.6) | 1-98 |  |
| WJ-IV Sound Awareness | 37.3 (25.1) | 0.1-97 |  |
| **Phonological Short-term Memory** |  |  |  |
| WJ-IV Nonword Repetition | 48.1 (25.2) | 1-99 |  |
| WJ-IV Memory for Words | 37.1 (25.5) | 0.3-99 |  |
| **Semantic Impact** |  |  |  |
| WJ-IV Sentence Repetition | 44.3 (24.7) | 2-94 |  |
| NEPSY-II Semantic Fluency | 76.7 (23.7) | 2-99.9 |  |
| **Rapid Naming** |  |  |  |
| WJ-IV Rapid Picture Naming | 30.2 (21.3) | 0.3-88 |  |
| **Expressive Language** |  |  |  |
| WJ-IV Oral Vocabulary | 51.6 (26.1) | 0.2-99.8 |  |
| **Registration & Working Memory** |  |  |  |
| WISC-IV Digit Span Forward | 35.4 (28.0) | 1-98 |  |
| WISC-IV Digit Span Backward | 34.0 (23.4) | 1-99 |  |
| **Processing Speed** |  |  |  |
| NEPSY-II Naming | 32.9 (27.4) | 0.5-99 | 2 |
| WISC-IV Symbol Search | 45.9 (25.4) | 1-98 | 1 |
| Children's Color Trails Test 1 | 37.4 (28.8) | 0.5-92 | 6 |
| WISC-IV Coding | 25.6 (24.4) | 0.5-99.5 |  |
| **Learning & Memory** |  |  |  |
| CVLT-C Trial 1 | 51.5 (29.5) | 2-99.5 |  |
| Rey-O 3-min Delayed Recall | 33.2 (29.8) | 0.5-98 | 1 |
| **Visuospatial & Visuomotor** |  |  |  |
| Judgement of Line Orientation | 40.7 (31.7) | 0.002-98 | 2 |
| Beery VMI | 33.8 (24.5) | 1-97 | 5 |
| **Executive Functioning** |  |  |  |
| NEPSY-II Inhibition | 39.1 (28.1) | 0.5-99.5 | 19 |
| NEPSY-II Switching | 40.8 (26.1) | 0.1-99 | 21 |
| NEPSY-II Initial Letter Fluency | 33.5 (27.3) | 0.5-99 |  |
| Children's Color Trails Test 2 | 28.1 (23.6) | 0.5-88 | 5 |
| **Mathematics** |  |  |  |
| WJ-IV Calculations | 23.2 (18.8) | 0.01-82 | 3 |

*Abbreviations* = Neuropsychological test scores are reported in both percentiles and z-scores; means (*M*), standard deviations (*SD*), ranges, and number of missing scores per task are reported. Abbreviations: WASI = Wechsler Abbreviated Scale of Intelligence (Wechsler, 1999); ROWPVT-4 = Receptive One-Word Picture Vocabulary - Fourth Edition (Martin & Brownell, 2011); TOWRE-2 = Test of Word Reading Efficiency – Second Edition (Torgesen et al., 2012); WJ-IV = Woodcock Johnson Fourth Edition – Test of Achievement (Schrack et al., 2014); NEPSY-II = NEPSY – Second Edition (Korkman et al., 2007); WISC-IV = Wechsler Intelligence Scale for Children – Fourth Edition Integrated (Wechsler et al., 2004); CVLT-C = California Verbal Learning Test – Children's Version (Delis et al., 1994); Rey-O 3-min Delayed Recall = Rey Osterrieth Complex Figure (Osterrieth, 1944; Rey, 1941); Judgement of Line Orientation = Judgement of Line Orientation – Form “H” (Benton et al., 1983); Beery VMI = Beery Buktenica Developmental Test of Visual-Motor Integration – Fourth Edition (Beery, 1997).

To further evaluate behavioral symptoms and identify ADHD, parents or legal guardians completed the Behavior Assessment System for Children—Second Edition (BASC-2; Kamphaus & Reynolds, 2007) or Third Edition (BASC-3; Kamphaus & Reynolds, 2015), along with the Vanderbilt ADHD Diagnostic Rating Scales—Parent and Teacher Reports (Wolraich et al., 2003).

Participants also completed 3T MRI scans, including structural, diffusion, and resting-state functional sequences.

### 2.3 Inclusion & Exclusion Criteria

The initial cohort consisted of 277 children assessed at the UCSF Dyslexia Center. A total of 130 were excluded for the following reasons: having a diagnosis of autism spectrum disorder (ASD); a reported history of prenatal alcohol exposure or global learning impairment; or a score below the 9th percentile on non-verbal reasoning and/or receptive vocabulary measures. Children were also excluded if they had clinically significant MRI findings.

Participants were included if the multidisciplinary team’s evaluation diagnosed developmental dyslexia based on the International Dyslexia Association’s definition (“Definition of Dyslexia”, 2002) and if they scored below the 20th percentile on at least one of the following measures: the Sight Word Efficiency or Phonemic Decoding Efficiency subtests of TOWRE-2 (Torgesen et al., 2012), or the Rate, Accuracy, or Comprehension subtests of GORT-5 (Wiederholt & Bryant, 2012). The 20th percentile threshold (below the low average range) was selected to increase sensitivity to reading impairment, as many children had received intensive reading intervention or tutoring that may have partially remediated their skills.

To be included, participants were also required to complete WASI Matrix Reasoning and ROWPVT-4, as well as at least 32 out of the 35 total standardized measures, corresponding to a minimum 90% task completion threshold. Importantly, children were not excluded on the basis of co-occurring ADHD, as it frequently co-occurs with dyslexia (Willcutt et al., 2010; Willcutt et al., 2005). Excluding these participants would reduce both the generalizability and the clinical relevance of the findings.

### 2.4 Statistical Analyses

We implemented a three-step analytic approach: (1) Hierarchical clustering to empirically group cognitive measures into data-driven clusters, (2) Latent profile analysis (LPA) to identify potential subgroups of children across cognitive domains, and (3) post-hoc group comparisons to evaluate outcomes and co-occurring conditions across identified profiles. In the LPA, age at testing, sex, school type, non-verbal reasoning (WASI Matrix Reasoning), and receptive vocabulary (ROWPVT-4) were included as covariates to account for demographic and IQ factors. Our approach allowed us to uncover latent subgroups of children and understand their distinct clinical and cognitive features.

### 2.4.1 Hierarchical Clustering of Tasks

To characterize distinct cognitive subdomains based on performance among the cohort of children with dyslexia, we conducted hierarchical clustering on percentile scores across 24 tasks from our assessment battery; these included (from Table 2) measures of phonological manipulation, phonological short-term memory, semantic impact, rapid naming, expressive language, registration and working memory, processing speed, learning and verbal memory, visuospatial and visuomotor skills, executive functioning, and mathematics. We excluded the tasks that directly assessed reading and spelling abilities (see single word reading, timed paragraph reading, and single word spelling sections in Table 2) as, by definition, these are known areas of difficulty in all children in the cohort.

The selection of these 24 measures was done by multidisciplinary consensus and considered the clinical relevance to neurodevelopmental disorders, insight into the differential diagnosis of dyslexia and related conditions, and feasibility of administration. We prioritized standardized tasks with normative data and tasks with minimal missing data across participants for the purposes of statistical robustness.

Hierarchical clustering was performed using Ward’s method with Euclidean distance, as this approach minimizes within-cluster variance and is well-suited for standardized data (Ward, 1963). Median imputation was used to account for missing scores in the dataset (1.90% missing). The cluster threshold was set at 70% of the maximum linkage distance, balancing between overly granular clustering and overly consolidated grouping as well as taking into consideration meaningfulness of cluster interpretation.

### 2.4.2 Latent Profile Analysis

Next, we performed a latent profile analysis (LPA) to distinguish distinct cognitive phenotypes among our cohort of children with dyslexia. We selected LPA due to its robustness in handling mixed data types, missing data, and utility in identifying unobserved subgroups in a population (Sinha et al., 2021). The LPA approach allowed us to maximize participant inclusion while maintaining accuracy of profiles by managing missing data points through maximum likelihood estimation, without over-introducing bias to the data distributions. This choice was crucial in increasing the statistical power and generalizability of findings. The analysis was completed with percentile scores from the 24 tasks mentioned above. Tasks were z-scored within subject to obtain relative strengths and weaknesses of each child. The following covariates were included as structural data (external factors that may influence the latent components): age at testing, sex, school type, WASI Matrix Reasoning, and ROWPVT-4. Bayesian Information Criterion (BIC) and Consistent Akaike Information Criterion (CAIC) were calculated to evaluate the optimal number of components (range: 2-10) for the LPA.

To assess the stability of the LPA results, we used a bootstrap resampling approach and measured alignment between bootstraps with the Adjusted Rand Index (ARI). We fit the latent profile model 1,000 times using bootstrapped samples with replacement, each containing 90% of the original dataset. We then computed pairwise ARI values between all possible pairs of bootstrap profile assignments to quantify their agreement. Cluster profiles were examined across each task for the average z-scored percentile (and standard error of the mean [SEM]) per cluster, as well as the original percentile scores before z-scoring. The LPA analysis and bootstrapping validation was repeated with scaled scores (z-scores generated from psychometric tables validated on large normative samples) which were also z-scored by each participant to obtain relative strengths and weaknesses of each child.

### 2.4.3 Post-hoc Comparisons Between Profiles

Post-hoc statistical comparisons between the two profiles were performed for variables that were entered in the LPA (Table S2) as well as additional reading and spelling measures (Table 3) and the frequency of co-occurring diagnosis between profiles (Figure 3). We corrected for multiple comparisons with false discovery rate (FDR) using the Benjamini & Hochberg procedure (Benjamini & Hochberg, 1995).

**Table 3.**
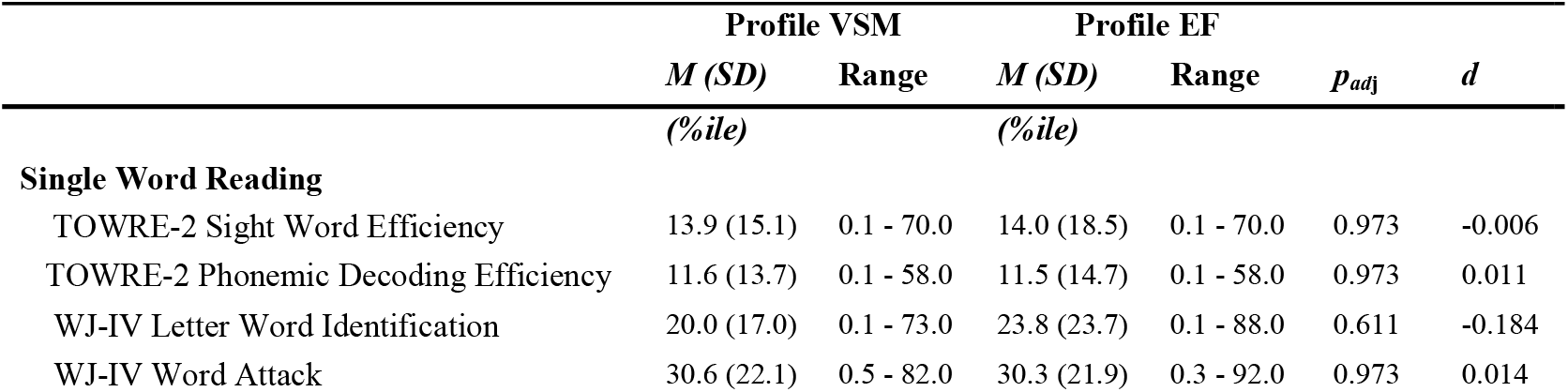
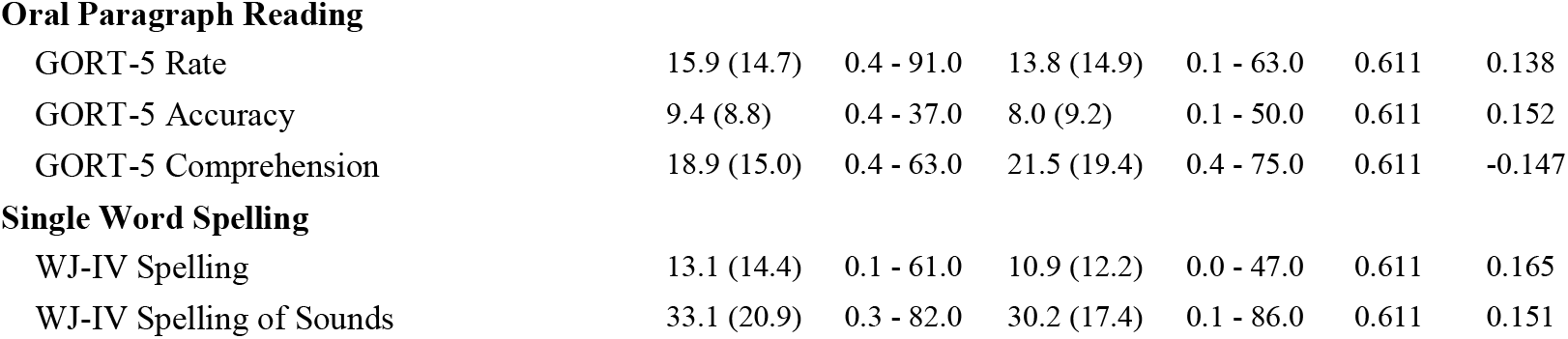
Performance on Reading and Spelling Measures by Profile.

| | Profile VSM | | Profile EF | | $p_{adj}$ | $d$ |
| --- | --- | --- | --- | --- | --- | --- |
|  | *M* (*SD*) | Range | *M* (*SD*) | Range |  |  |
|  | (%ile) |  | (%ile) |  |  |  |
| **Single Word Reading** |  |  |  |  |  |  |
| TOWRE-2 Sight Word Efficiency | 13.9 (15.1) | 0.1 - 70.0 | 14.0 (18.5) | 0.1 - 70.0 | 0.973 | -0.006 |
| TOWRE-2 Phonemic Decoding Efficiency | 11.6 (13.7) | 0.1 - 58.0 | 11.5 (14.7) | 0.1 - 58.0 | 0.973 | 0.011 |
| WJ-IV Letter Word Identification | 20.0 (17.0) | 0.1 - 73.0 | 23.8 (23.7) | 0.1 - 88.0 | 0.611 | -0.184 |
| WJ-IV Word Attack | 30.6 (22.1) | 0.5 - 82.0 | 30.3 (21.9) | 0.3 - 92.0 | 0.973 | 0.014 |
| **Oral Paragraph Reading** |  |  |  |  |  |  |
| GORT-5 Rate | 15.9 (14.7) | 0.4 - 91.0 | 13.8 (14.9) | 0.1 - 63.0 | 0.611 | 0.138 |
| GORT-5 Accuracy | 9.4 (8.8) | 0.4 - 37.0 | 8.0 (9.2) | 0.1 - 50.0 | 0.611 | 0.152 |
| GORT-5 Comprehension | 18.9 (15.0) | 0.4 - 63.0 | 21.5 (19.4) | 0.4 - 75.0 | 0.611 | -0.147 |
| **Single Word Spelling** |  |  |  |  |  |  |
| WJ-IV Spelling | 13.1 (14.4) | 0.1 - 61.0 | 10.9 (12.2) | 0.0 - 47.0 | 0.611 | 0.165 |
| WJ-IV Spelling of Sounds | 33.1 (20.9) | 0.3 - 82.0 | 30.2 (17.4) | 0.1 - 86.0 | 0.611 | 0.151 |

## 3. Results

### 3.1 Neuropsychological Assessment Results

The mean percentile scores across the neuropsychological test battery reflected a broad range of abilities in our cohort. Table 2 presents a detailed summary of these results.

### 3.2 Hierarchical Clustering of Tasks

Hierarchical clustering of neuropsychological tasks identified four distinct partially independent clusters (correlations between r = 0.16-0.48; Figure 1). For interpretation of clusters, we assigned each a cognitive domain label based on the commonalities among the tasks: (1) Processing Speed & Executive Function, (2) Visuospatial & Mathematical Reasoning, (3) Phonological Manipulation, (4) Verbal Short-term Memory & Word Retrieval.

**Figure 1.**
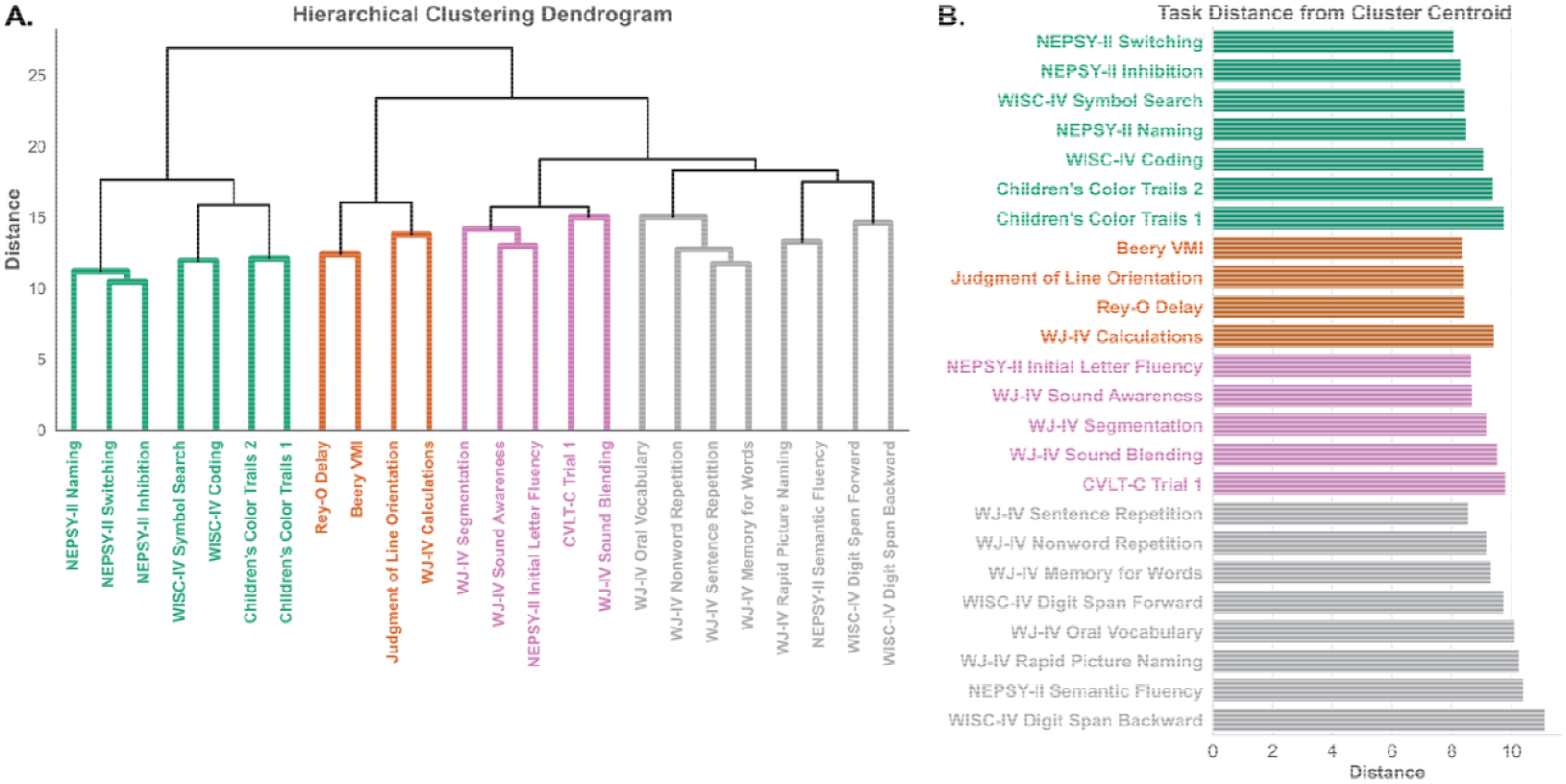
Hierarchical clustering analysis of neuropsychological measures. **A**. Dendrogram showing hierarchical relationships between tasks, with height indicating linkage Euclidian distance between clusters. **B**. Distance from respective cluster centroid for 24 measures, grouped into four clusters.

### 3.3 Latent Profile Analysis

Model fit indices, including the Bayesian Information Criterion (BIC) and Corrected Akaike Information Criterion (CAIC), indicated that a two-profile solution best fit the data. Accordingly, the LPA was conducted with two profiles (Figure 2). Children in Profile VSM (n = 75) showed relative difficulties in verbal short-term memory and word retrieval, but average performance in processing speed and executive functioning. In contrast, children in Profile EF (n = 72) exhibited relative weaknesses in processing speed and executive functioning, while performing on par in verbal short-term memory and word retrieval. Thereafter, we will refer to Profile VSM as VSM (verbal short-term memory) and Profile EF as EF (executive functioning).

**Figure 2.**
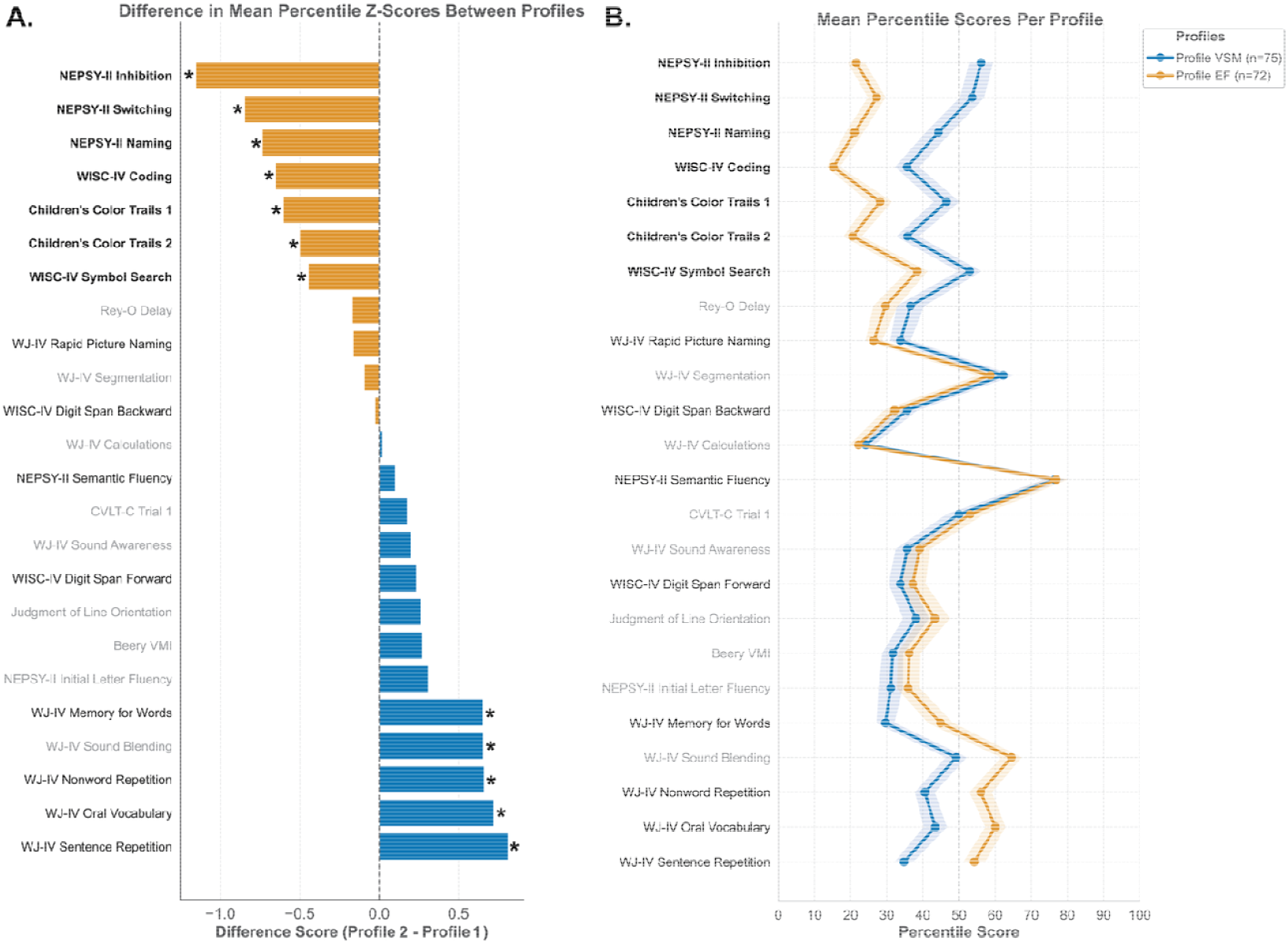
**A. Difference in mean percentile z-score between children in Profile VSM versus Profile EF**.Profile VSM mean scores were subtracted from Profile EF to create a difference score representing performance disparity between groups. Negative values indicate lower performance in Profile EF (relative weakness in processing speed and executive functioning). Positive values indicate lower performance in Profile VSM (relative weakness in verbal short-term memory and word retrieval). Bold task labels indicate the processing speed and executive function tasks, normal weight task labels indicate verbal short-term memory and word retrieval tasks, and grey task labels indicate tasks from the remaining clusters. Asterisks indicate significant t-tests after FDR correction for post-hoc comparisons between profiles. **B. Mean percentile scores grouped by profile**. The lines indicate mean task performance per profile, with the line shadings representing standard error of the mean (SEM). Tasks are ordered according to differences in percentile z-scores.

**Figure 3.**
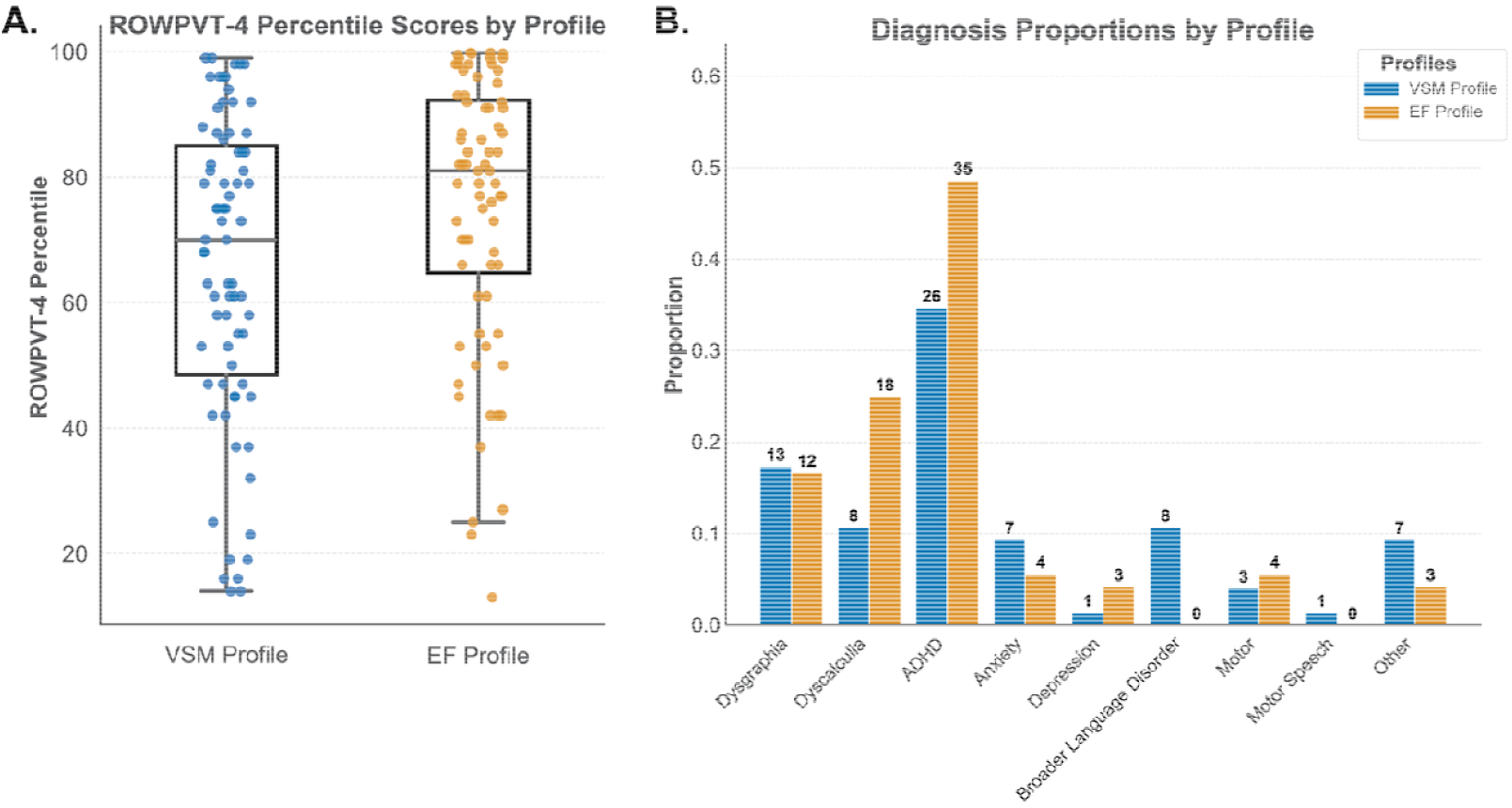
**A. Boxplots for ROWPVT-4 percentile scores**.Plot indicates first quartile, median, and third quartile; whiskers show first and third quartile ±1.5*interquartile range. **B. Proportion of co-occurring diagnoses per profile**. Diagnosis of “Other” was included: childhood apraxia of speech, impulse control concerns, articulation concerns, speech sound disorder, possible language disorder.

Both groups demonstrated clear relative strengths in semantic fluency with more than half of children in each group performing above the 75^th^ percentile (Profile VSM = 54.7%; Profile EF = 55.6% respectively). Children in both profiles also demonstrated relative weaknesses in visuospatial skills with more than a third from each profile performing below the 25^th^ percentile in Judgement of Line Orientation (Profile VSM = 38.7%; Profile EF = 38.9%), and mathematical processing with more than half performing below the 25^th^ percentile in WJ-IV Calculations (56.0% and 65.3% respectively; see Figure 2b). Refer to Supplemental Table S2 for a summary of performance on the 24 tasks per profile.

As displayed in Figure 2a, significant differences between profiles after FDR correction (all *p*_adj_<0.001, see Table S2) were found in all seven tasks of the Processing Speed & Executive Function cluster (NEPSY-II Inhibition, NEPSY-II Switching, NEPSY-II Naming, WISC-IV Coding, Children’s Color Trails 2, Children’s Color Trails 1, WISC-IV Symbol Search), four tasks in the Verbal Short-term Memory & Word Retrieval cluster (WJ-IV Sentence Repetition, WJ-IV Oral Vocabulary, WJ-IV Nonword Repetition, WJ-IV Memory for Words), and one task in the Phonological Manipulation cluster (WJ-IV Sound Blending).

The bootstrap resampling approach to quantify stability resulted in an average Adjusted Rand Index of 0.648 (*SD* = 0.210), which indicated significant agreement between latent profile solutions. Importantly, performance between classes was not significantly different in any measure of reading or spelling (see Table S2), suggesting that the subtypes identified are independent from variations in reading or spelling ability alone. Children in Profile EF presented with a higher proportion of co-occurring ADHD diagnoses, but group differences in the number of children with or without ADHD was not significant between the two profiles (VSM Profile: N=26 [34.7%] ADHD, EF Profile: N=35 [48.6%], *χ*^2^ = 2.40; *p*_adj_= 0.365). There was a statistically higher incidence of dyscalculia in Profile EF which did not remain significant after FDR correction (dyscalculia: ⍰^2^ = 4.25; *p*_adj_ = 0.117).

There was no significant difference in the number of total co-occurring diagnoses per child between the two profiles as shown by a Mann-Whitney U test (U=2409.50; *p*=0.235). Children in both groups had an average of one co-occurring diagnosis in addition to dyslexia. Among model covariates, there was a significant difference in receptive vocabulary (ROWPVT-4) between the two profiles (*t*=-2.447; *p*=0.016) before FDR correction, however the p-value became insignificant after correction (*p*_adj_=0.093). There was also no significant difference in age (*t*=1.359; *p*=0.352), sex (*χ*^2^=3.065; *p*_adj_=0.240), school type (*χ*^2^=0.120; *p*_adj_=0.942), or handedness (*χ*^2^=0.788; *p*_adj_=0.480) between the two profiles.

## 4. Discussion

Developmental dyslexia is a common neurodevelopmental condition characterized by persistent reading difficulties. Dyslexia is increasingly recognized as a multifactorial disorder, and interventions that target only phonological or reading-specific symptoms are often insufficient. This underscores the need for comprehensive assessment and multidisciplinary consensus to capture the diverse underlying cognitive factors beyond reading-related symptoms. Despite its prevalence and significant impact on life outcomes (Georgiou et al., 2024; Wilmot et al., 2023), large-scale studies incorporating such multidisciplinary evaluations remain scarce. Our study aimed to identify distinct cognitive subtypes within dyslexia using a data-driven clustering approach. First, we grouped the assessment tasks into four cognitive domains: Processing Speed & Executive Function, Visuospatial & Mathematical Reasoning, Phonological Manipulation, and Verbal Short-term Memory & Word Retrieval. We then clustered over 140 children with persistent dyslexia based on their performance across the entire neuropsychological battery, which revealed two broader cognitive profiles. The cognitive strengths and weaknesses defining each profile were mapped onto the four previously delineated domains, although both groups displayed comparable levels of reading and spelling impairment. These findings move beyond traditional phonological models of dyslexia, suggesting that children with similar reading difficulties can present with divergent constellations of cognitive challenges. Importantly, they suggest that conventional literacy assessments, which focus predominantly on phonological processing, orthographic decoding, and rapid naming, may fail to capture critical dimensions of the cognitive profile that contribute to reading impairment (Ozernov-Palchik et al., 2016).

Children in the first profile exhibited relative weaknesses in tests that probed verbal short-term memory and word retrieval. Specifically, they performed poorly on WJ-IV Sentence Repetition, Nonword Repetition, and Memory for Words—tasks that depend on efficient use of the phonological loop 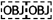. Previous work has shown that sentence repetition is a predictor of broader language impairments, conceptualized as a measure of integrity of language processing systems including speech perception and vocabulary knowledge (Klem et al., 2015; Potter & Lombardi, 1990) and often indicate 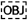(DLD;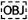

Additionally, children showed weaknesses in WJ-IV Oral Vocabulary which relies on word retrieval and expressive language abilities. Weak performance on WJ-IV Sound Blending, probing the ability to combine phonemes to form words, further indicates the shared reliance on phonological awareness. These findings underscore the role of verbal awareness and short-term memory in supporting reading acquisition and comprehension, and broader language outcomes (Brady, 1986; Majerus & Cowan, 2016). Numerous studies have highlighted that verbal short-term memory deficits are prevalent in dyslexia and predict single-word reading and reading comprehension (Gathercole, 2006; Gathercole & Baddeley, 1989; Swanson & Siegel, 2001). Notably, although many children in the verbal short-term memory profile exhibited difficulties consistent with DLD, only a relatively small proportion were formally diagnosed. This discrepancy may reflect differences in clinical identification (i.e., academic, neuropsychological, medical or speech pathology) or the language assessment administered (Bishop, 2017). Previous literature suggests that nonword repetition and sentence repetition are among the most sensitive markers for DLD, underscoring the need for comprehensive language assessment in children with dyslexia (Conti-Ramsden et al., 2001). Further work is necessary to evaluate the interplay between auditory short-term memory difficulties in dyslexia, ADHD, and DLD.

Children in the second profile were characterized by relative difficulties in executive functioning and processing speed. More specifically, this profile showed impairment in tasks such as NEPSY-II Inhibition and Switching, which require the ability to control impulses and shift attention efficiently between tasks (De Rom et al., 2023). These difficulties were corroborated by lower performance on WISC-IV Coding and Symbol Search, as well as Children’s Color Trails 1 and 2, which collectively underscore the marked deficits in processing speed and cognitive control (Diamond, 2013). Executive functions including working memory, cognitive flexibility, and inhibitory control, are increasingly recognized as essential for successful reading (Friedman & Robbins, 2022; Lonergan et al., 2019).

While previous models have emphasized deficits in phonological processing, rapid automatized naming, and orthography (King et al., 2007; O’Brien et al., 2012; Peterson et al., 2013; Sprenger-Charolles et al., 2000; Wolf & Bowers, 1999) our study demonstrates that dyslexia is a multifaceted condition, with distinct cognitive aspects being associated with reading disorder (Alt et al., 2022; Kramer et al., 2000; Majerus & Cowan, 2016). Despite the observation of a higher incidence of ADHD in the PS-EF profile, the absence of statistically significant differences in ADHD diagnoses between profiles suggests that the identified cognitive weaknesses may operate independently of ADHD (Al Dahhan et al., 2022). This finding held true for all other co-occurring conditions among our sample. Taken together, our results challenge reliance on reading assessments alone and align with the multiple-deficit model, which posits that reading difficulties arise from the interaction of multiple cognitive risk factors (Pennington et al., 2012; van Bergen et al., 2014). This model is supported by accumulating evidence that children with similar reading impairments may have different constellations of underlying cognitive weaknesses, including deficits in executive function, processing speed, and verbal memory.

Several features of this study should be acknowledged. First, our sample was clinically biased, as it was drawn primarily from intervention-based learning environments, which may limit the generalizability of the findings to the broader population of children with dyslexia (Snowling, 2013). Furthermore, the dyslexia intervention curriculum in our partner schools focuses heavily on phonological training, likely boosting participants’ phonological processing skills and minimizing differences between profiles in this domain. It is possible that, in the absence of such intervention, we would have observed a group characterized primarily by phonological deficits—similar to what has been reported in studies of less-remediated populations (van Ermingen-Marbach et al., 2013). Conversely, this uniformity in phonological skills within our cohort created a valuable opportunity to detect subtypes distinguished by non-phonological weaknesses, such as deficits in executive functioning or processing speed. This aligns with the perspective that dyslexia encompasses a broader spectrum of cognitive difficulties beyond phonological deficits (Snowling & Hulme, 2012a).

Future research should aim to replicate these findings in larger, more diverse cohorts and explore longitudinal trajectories to determine how these cognitive profiles evolve throughout development and with targeted intervention. The cross-sectional design of the present study precludes conclusions about the developmental stability of these profiles or their responsiveness to intervention over time. Moreover, integrating neuroimaging data could provide insights into the neural mechanisms that differentiate these subtypes, potentially guiding the refinement of both theoretical models and practical intervention strategies. Weaknesses in processing speed and executive function in Profile EF may reflect differences or dysfunction in the fronto-parietal network, which coordinates executive processes through interactions between prefrontal and parietal regions (Marek & Dosenbach, 2018; Ptak, 2011). Conversely, difficulties with verbal short-term memory and word retrieval as seen in the other emergent profile in our study may point to potential disruptions in the left perisylvian network, including the inferior frontal gyrus, superior temporal gyrus, and the arcuate fasciculus connecting these regions, which are critical for phonological processing, verbal memory, and language production (Catani et al., 2005; Koenigs et al., 2011; Riès et al., 2017).

Future efforts should translate these findings into scalable assessment batteries. While brief, easy-to-administer dyslexia screeners might be sufficient for determining risk of dyslexia, a multi-domain neuropsychological testing battery is necessary for a formal diagnosis and painting the distinct cognitive profile of the child (Sanfilippo et al., 2020). Administering a fixed comprehensive assessment allows direct comparison of measures across varying cognitive profiles, offering several advantages over a flexible battery approach (Casaletto & Heaton, 2017). However, implementing such batteries at scale presents significant financial and logistical challenges, including the need for substantial clinician training, increased testing time, and careful coordination with school resources—factors that can limit feasibility in typical educational or clinical settings (Lezak et al., 2012).

Addressing the mentioned barriers will be crucial for developing more accurate assessment protocols that enable tailored interventions based on each child’s needs (Mather & Schneider, 2023). Current intervention strategies typically adopt a “one-size-fits-all” approach focused on phonological training (Wanzek et al., 2018). However, the distinct behavioral profiles identified here could explain why traditional phonological interventions may not fully address reading difficulties in all children with dyslexia. Targeted interventions (such as executive-function training for children with EF impairments or explicit memory-strategy instruction for those with verbal short-term-memory weaknesses) may therefore prove more effective.

In summary, this study underscores the cognitive heterogeneity inherent in reading impairments and highlights the potential of data-driven methods to uncover distinct cognitive profiles. By moving beyond traditional reading-centric assessments, our findings advocate for a more nuanced, individualized approach to both diagnosis and intervention. This approach not only advances theoretical understanding of dyslexia but also holds practical promise for improving educational and clinical outcomes. Clinically, our results underscore the need for comprehensive, multidomain assessment in children with persistent reading difficulties, as conventional screeners can miss meaningful cognitive differences that should guide intervention (Sanfilippo et al., 2020). By using each child’s unique cognitive profile, educators and clinicians can tailor interventions and, ultimately, optimize reading outcomes.

## Author Contribution

Conceptualization: PPC, MK, MLGT.

Data Curation: MK, RB, EC, DVMM, SI

Formal Analysis: MK

Funding Acquisition: PPC, MLGT

Investigation: RB, EC, DVMM, SI, JDL, BLT, CP, ZM, MLGT

Methodology: PPC, MK, JV, MLM, MLGT

Project Administration: SI, CP, MLGT

Resources: PPC, MLGT

Software: MK, PPC,

Supervision: PPC, MLGT

Validation: MK

Visualization: MK

Writing - Original Draft Preparation: MK, PPC

Writing - Review & Editing: MK, PPC, NB, SI, JS, CP, MLGT

### Acknowledgements

We are deeply grateful to the families and children who generously participated in this study and whose dedication made this research possible. We thank our partnering schools, Charles Armstrong School and Chartwell School, for their commitment and support in facilitating recruitment and assessment. Special thanks to our research coordinator, whose tireless efforts ensured seamless study organization, data collection, and communication with families. We also acknowledge the outstanding support from UCSF Information Technology (IT), whose expertise was vital in the implementation, management, and secure storage of our large-scale datasets. Finally, we are indebted to the multidisciplinary team at the UCSF Dyslexia Center for their clinical insight, collaboration, and steadfast dedication to advancing understanding and care for children with learning differences.

## Code Availability

All data preparation and statistical analyses were conducted using Python 3.11.9. Hierarchical clustering was performed using SciPy package version 1.11.4 (Gommers et al., 2023). Latent Profile Analysis was conducted using Stepmix package version 2.2.1 (Morin et al., 2023). All the scripts supporting the analyses are available on our GitHub repository (https://github.com/MathCognitionUCSF/dyslexia_phenotype_clusters).

## Data availability statement

Academic, not-for-profit investigators can request subjects, tissue and laboratory specimens, archived and imaging data, technological tools or video clips of behaviors for professional education and research for research studies from the UCSF Memory and Aging Center, but you must have Institutional Review Board approval from the UCSF Human Research Protection Program (HRPP). For more information, see https://memory.ucsf.edu/research-trials/professional/open-science.

## Gender citation balance

We examined the gender balance of the works cited in our reference list by inferring the likely gender of first and last authors from their first names, using publicly available publication metadata. Authors were coded as woman or man based on common name usage. Cases in which the author was an organization, only initials were available, or the first name was highly ambiguous were coded as unknown and excluded from proportion calculations. Of the 91 references in our bibliography, we were able to assign a binary gender to both first and last authors for 84 references. Among these, 55.2% of first authors and 37.6% of last authors were inferred to be women. Considering first and last authors jointly, 26.2% of citations had a woman as both first and last author, 29.8% had a woman first author and man last author, 10.7% had a man first author and woman last author, and 33.3% had a man as both first and last author.

---

**Table 1.**
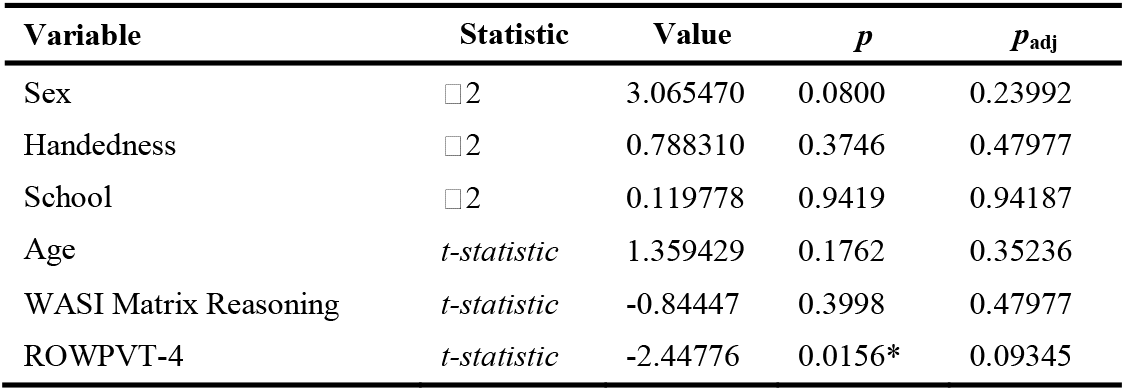
Summary of Statistical Tests for Group Differences Across Demographic and Cognitive
Variables.

**Table 2.**
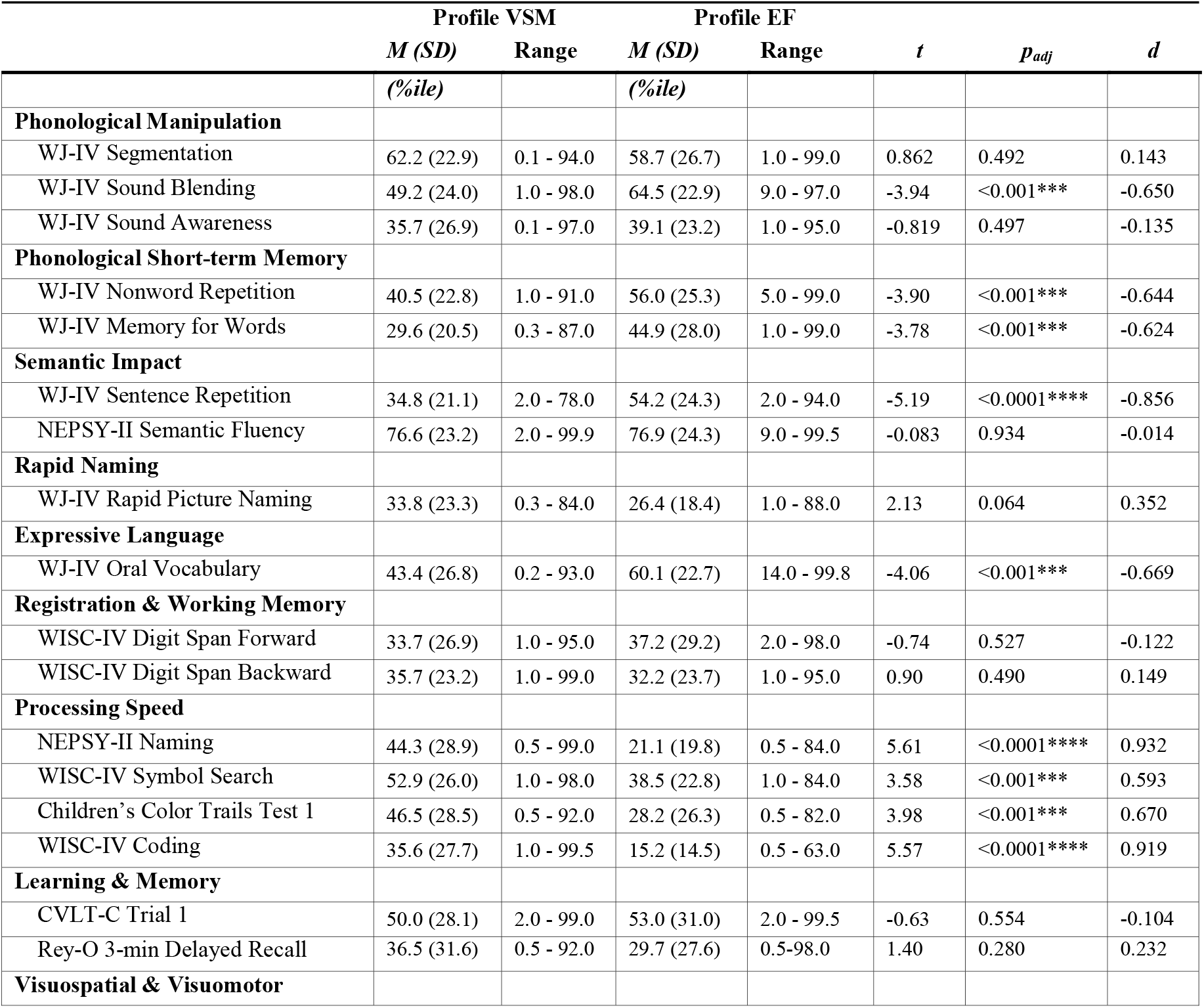
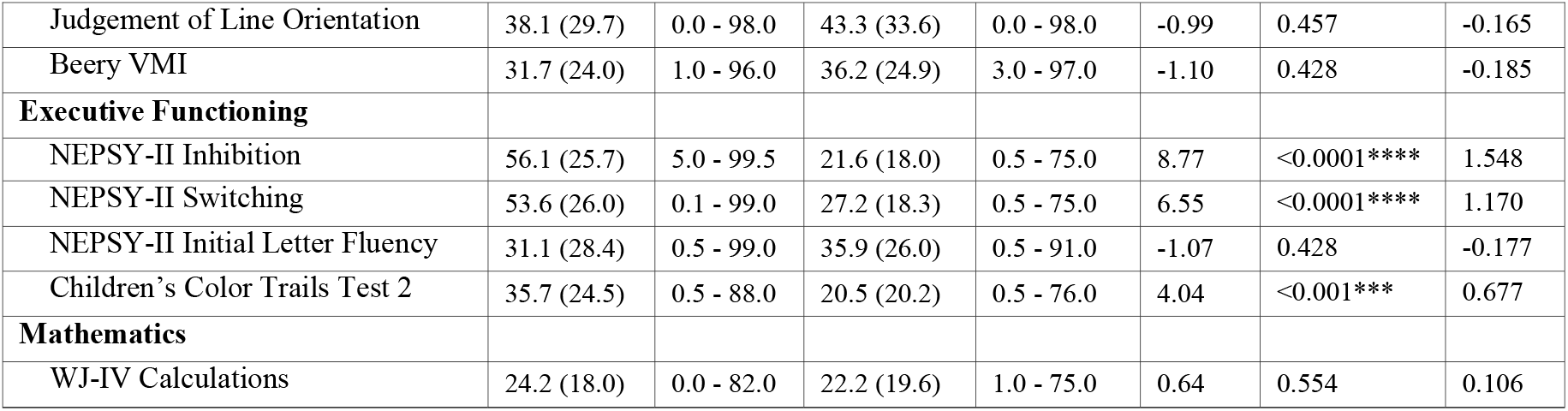
Performance on LPA Cognitive Measures by Profile.

## Notes

### Competing Interest Statement

The authors have declared no competing interest.

### Funding

- Charles and Helen Schwab Foundation

